# *Ex vivo* SIM-AFM measurements reveal the spatial correlation of stiffness and molecular distributions in 3D living tissue

**DOI:** 10.1101/2024.05.27.595975

**Authors:** Itsuki Shioka, Ritsuko Morita, Rei Yagasaki, Duligengaowa Wuergezhen, Tadahiro Yamashita, Hironobu Fujiwara, Satoru Okuda

**Affiliations:** Graduate School of Frontier Science Initiative, Kanazawa University, Kanazawa 920-1192, Japan; Graduate School of Frontier Biosciences, Osaka University, Osaka 565-0871, Japan; Nano Life Science Institute, Kakuma-machi, Kanazawa 920-1192, Japan; Laboratory for Tissue Microenvironment, RIKEN Center for Biosystems Dynamics Research (BDR), Kobe 650-0047, Japan; Department of System Design Engineering, Faculty of Science and Technology, Keio University, Yokohama 223-8522, Japan

## Abstract

Living tissues each exhibit a distinct stiffness, which provides cells with key environmental cues that regulate their behaviors. Despite this significance, our understanding of the spatiotemporal dynamics and the biological roles of stiffness in three-dimensional tissues is currently limited due to a lack of appropriate measurement techniques. To address this issue, we propose a new method combining upright structured illumination microscopy (USIM) and atomic force microscopy (AFM) to obtain precisely coordinated stiffness maps and biomolecular fluorescence images of thick living tissue slices. Using mouse embryonic skin as a representative tissue with mechanically heterogeneous structures inside, we validate the measurement principle of USIM-AFM. Live measurement of tissue stiffness distributions revealed the highly heterogeneous mechanical nature of embryonic skin as well as the role of collagens in maintaining its integrity. Furthermore, quantitative comparisons of stiffness distributions of preserved tissue samples unveiled the distinct impacts of conventional tissue preservation techniques on the tissue stiffness pattern. This series of experiments highlights the importance of live mechanical testing of tissue-scale samples. Our USIM-AFM technique provides a new methodology to reveal the dynamic nature of tissue stiffness and its correlation with biomolecular distributions in live tissues and thus could serve as a technical basis for exploring tissue-scale mechanobiology.

**Graphical Abstract:** 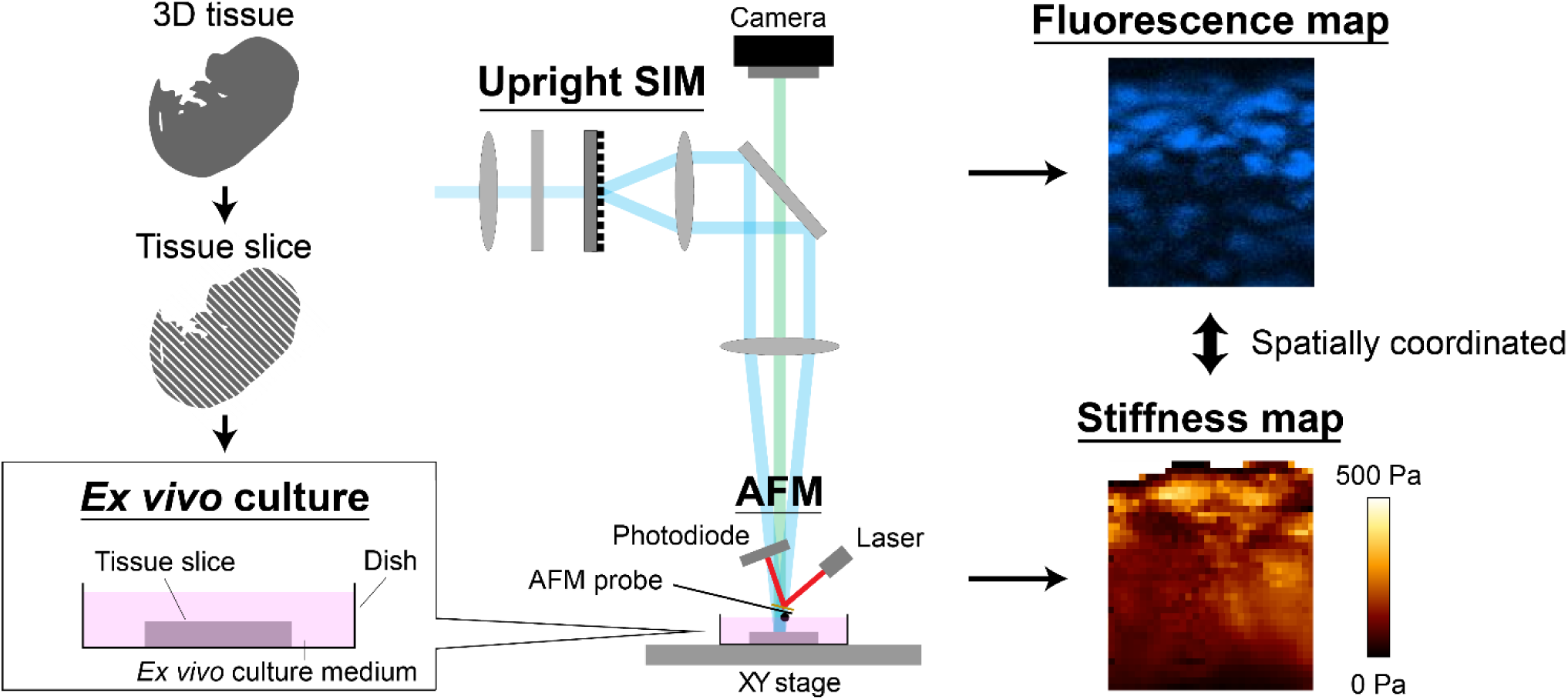

## 1. Introduction

Tissue stiffness exhibits heterogeneity throughout the body, varying according to tissue type, developmental stage, and disease presence. For example, the brain [1,2] and lungs [1] display low stiffness, whereas bones [1,3] and skeletal muscles [1,4,5] exhibit high stiffness; additionally, tumors are typically stiffer than their respective normal tissues [6]. Moreover, tissue stiffness changes dynamically during development processes [7] and cancer progression [8,9]. This spatial and temporal variation in tissue stiffness plays various roles in biological systems. For example, cells respond to surrounding stiffness, affecting cell deformation and survival [10], differentiation [1,3], proliferation [11,12], and migration [13–19]. Such mechanical aspects of the cell culture environment have been actively implemented into cutting-edge biomaterial designs, *e.g.*, four-dimensional controllable hydrogels [20–23], aiming for spatiotemporal control of mechanical and chemical properties of the cell culture environment. However, our current understanding of the spatiotemporal pattern of stiffness and its role in regulating cell behaviors in living three-dimensional (3D) tissues remains limited due to the lack of a method to reveal the spatial heterogeneity of tissue stiffness and its link to the biological properties of cells positioned deep inside the target. There is thus an increasing need in both biological and bioengineering contexts for a new measurement method that simultaneously obtains the local tissue stiffness and biomolecular distributions within living 3D tissues.

Obtaining a tissue stiffness map at the micron scale is technically challenging due to the limited access of a mechanical probe to regions of interest positioned deep inside the tissue. To address this issue, non-invasive measurement techniques such as ultrasound elastography, magnetic resonance elastography (MRE), and Brillouin microscopy have been utilized [24]. Although these techniques provide contactless methods to obtain a stiffness map inside a living body, ultrasound elastography and MRE offer spatial resolutions limited to several hundred micrometers [24–26]. Brillouin microscopy, an advanced form of optics-based elastography, offers a spatial resolution of several hundred nanometers. However, the direct correlation of the obtained values with the target stiffness remains under debate, and the applicability of Brillouin microscopy for mechanically heterogeneous targets needs to be carefully validated from the viewpoint of mechanics [27]. However, there has been considerable progress in the application of atomic force microscopy (AFM), a form of contact-based scanning probe microscopy, for cellular measurements correlating the stiffness and cellular dynamics [28–33]. AFM is one of the most reliable tools to directly assess the target stiffness based upon a force-distance relationship, regardless of whether the target is linearly elastic. Additionally, AFM offers flexible tuning of the spatial resolution, ranging from 10 nm to 200 μm. Because of these advantages, AFM measurement has been utilized to test the mechanical properties of various biological targets, even at the subcellular scale, including the cytoskeleton [14,34,35], focal adhesions [36], nuclear surfaces [37], and cell surfaces [31–33,38]. Its use has recently been extended to tissue samples [7,39–42] including sliced tissue sections [41,43,44]; pioneering studies have obtained the statistical distribution of stiffness values in live embryonic brains and organoids [7,41,45]. However, recent reports further clarified that preservation, or even the transport process of the sample critically affects the stiffness values measured by AFM [41,43,46–49], highlighting the significance of conducting live and intact measurements at the tissue scale. Although AFM is a promising technique for mechanical testing of biological samples with fine spatial resolution, there is still room for improvement to extend its application to tissue-scale samples to pursue biological questions.

To explore tissue-scale cellular mechanics and mechanobiology, we herein developed a new AFM-based measurement system that reveals the spatial distribution of the local tissue stiffness arising from both intracellular and intercellular structures. There are two major requirements. First, the tissue sample needs to be sufficiently thick, *i.e.*, several hundred micrometers, so that the measured stiffness values reflect the genuine local stiffness of the continuous tissue. Second, a fluorescence microscope that can observe the top surface of the target, where AFM characterizes the mechanical properties, needs to be thoroughly integrated with the AFM measurement system. This requirement is practically indispensable for finding subcellular structures within optically opaque tissue and obtaining a fluorescence image aligned precisely to the stiffness map at the subcellular scale. However, these two requirements usually compete. A thicker tissue sample increases the difficulty of optically observing the top surface, the measurement field of AFM, using a common inverted microscope. To solve this dilemma, we propose a unique setup combining AFM and upright structured illumination microscopy (USIM). USIM provides a fluorescence image of the AFM measurement field; such imaging cannot be accomplished using conventional inverted fluorescence microscopes. The combined setup thus yields a paired stiffness map and fluorescence image of a thick tissue surface, guaranteeing their spatial correspondence at the micron scale. This setup even allows us to conduct spatiotemporal tracking of the stiffness changes in live 3D tissue together with precise correlation to the optical image when combined with an *ex vivo* slice culture chamber. Such live imaging could reveal the dynamic response of live tissue stiffness, which is lost with conventional preservation processes.

To demonstrate the measurement principle of USIM integrated with AFM (USIM-AFM), we analyzed the stiffness distribution of mouse embryonic skin consisting of a layered epidermis and mesenchyme with different cell densities and structures. The heterogeneous mechanical properties are currently considered to play substantial roles in regulating the cellular proliferation and differentiation that maintain the integrity of the skin [1,3,11,12]. To the best of our knowledge, there are no reports directly measuring the absolute stiffness map within live embryonic skin due to the lack of a measurement method. Utilizing the USIM-AFM system equipped with an *ex vivo* slice culture chamber, we reveal the stiffness distribution of such a complex tissue under a microscope. We first demonstrate the advantage of using USIM to distinguish the heterogeneous tissue structure of a live sample set in the AFM measurement field. Next, we show the importance of live tissue sectioning to determine the actual mechanical properties of 3D tissues by comparing the stiffness map detected on an intact skin surface with that detected on a sliced tissue section. We then demonstrate time-lapse imaging of the stiffness and nuclear distributions within a live tissue sample under drug treatment; such measurements are not practical with conventional methods. Finally, we reveal the importance of live stiffness measurement by comparing stiffness maps of live and differently preserved samples. Quantitative comparison of the stiffness distribution seen in the live and preserved tissues reveals different impacts of preservation processes on the absolute stiffness values as well as the texture of the stiffness map within the tissue. Our new methodology, USIM-AFM, sheds light on the value of live stiffness measurement of tissue-scale samples and thus serves as a new technical basis to pursue the mechanobiology of live tissues.

## 2. Materials and methods

### 2.1 Measurement system combining USIM, AFM, and *ex vivo* slice culture

An upright structured illumination microscope was integrated with an atomic force microscope to obtain a stiffness map together with a precisely aligned immunofluorescence image of sliced tissue (Fig. 1). The upright structured illumination microscope provides an optical image of the top surface of the tissue sample, whereas the atomic force microscope characterizes the mechanical properties, regardless of the sample thickness. An *ex vivo* slice culture chamber that maintained the viability of sliced tissue sections was also included on the microscope stage. Technically, an atomic force microscope head with a cantilever was positioned on the motorized stage of the atomic force microscope (Bruker, NanoWizard 4 Bioscience AFM JPK). The atomic force microscope head and stage were placed on the stage of a structured illumination microscope (Zeiss, Axiozoom) with a 0.5× objective (0.125 NA/114 mm WD) and a 1.0× objective (0.25 NA/56 mm WD) equipped with a CMOS camera (ZEISS, Axiocam 712 mono). A temperature-controlled chamber was installed on the atomic force microscope stage, and tissue slices were cultured *ex vivo* for measurement.

**Fig. 1.**
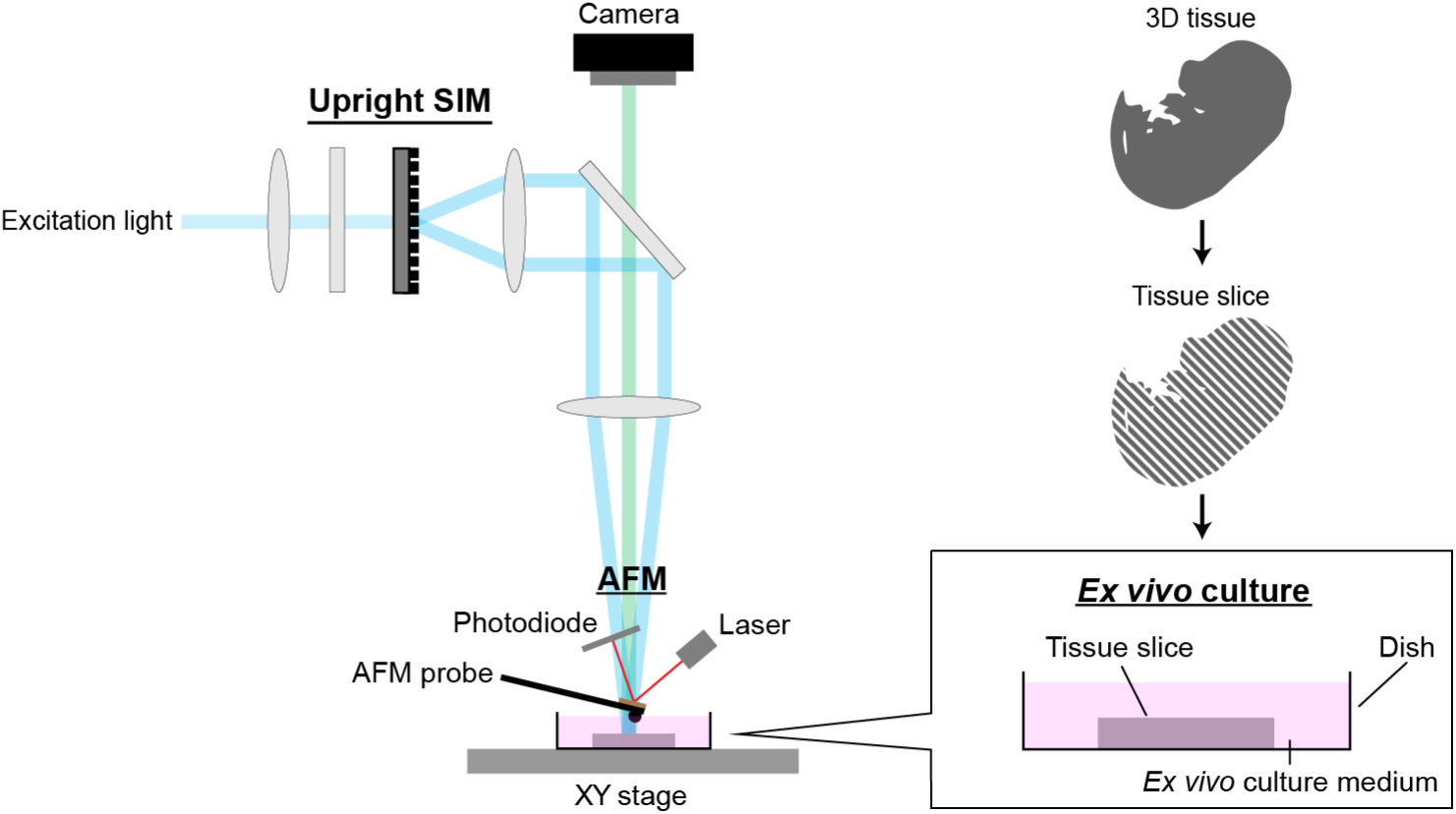
Schematic of the technique for simultaneous acquisition of stiffness and fluorescence maps at the same focal plane in living tissues. This technique integrates atomic force microscopy (AFM), upright fluorescence structured illumination microscopy (USIM), and *ex vivo* slice culture. An atomic force microscope head is mounted on a motorized stage, positioned between the upright fluorescence structured illumination microscope stage and the objective lens. The tissue slice is prepared, placed on the motorized stage, and maintained under *ex vivo* culture conditions throughout the measurement process.

### 2.2 Animals

The animal protocols for mice, including breeding and experiments, were approved by Kanazawa University and performed according to the guidelines of the Science Council of Japan. Pregnant ICR mice (Japn SLC, Inc.) were utilized for experiments within 2 days. Noon on the day that the vaginal plug was detected was defined as embryonic day 0.5 (E0.5). E14.5 mice were used in this study.

### 2.3 Preparation of living samples

Slice and surface samples of dorsal skin were prepared from E14.5 mice. First, embryos were extracted from the uterus in ice-cold Hank’s balanced salt solution (HBSS). The dorsal portion was cut from an embryo (along the red dotted line in Fig. 2**a**) and was collected in ice-cold HBSS. The skin was peeled from the dorsal portion using needles (Fig. 2**b**). The mesenchymal side of the skin was attached to the cellulose membrane (Fig. 2**c**). The following procedures were separately performed to prepare i) surface samples and ii) slice samples. The composition of the *ex vivo* culture medium used in the following procedures has been described in a previous study [50]. The medium consisted of Dulbecco’s modified Eagle’s medium and an F-12 nutrient mixture (Wako) supplemented with 20% fetal bovine serum (GIBCO), 1% penicillin-streptomycin (Wako), 2 mM GlutaMAX supplement (GIBCO), and 100 μg/ml ascorbic acid (Wako). The *ex vivo* culture medium was kept at 4 °C until use. To prepare surface samples (i), the skin attached to the cellulose membrane was stained with a 1:100 dilution of Hoechst 33342 (Wako, 34607951) in the *ex vivo* medium for 1 hour at 37 °C. The cellulose membrane with the skin was attached to a dish (Fig. 2**e**). To prepare slice samples (ii), the skin attached to the cellulose membrane was encapsulated in 4% agarose/phosphate-buffered saline (PBS) on the membrane to comprise a block including agarose gel, skin, and the membrane. The agarose gel was completely solidified in an ice bath. The block was cut with two parallel razor blades using a custom slicer to create a flat slice with the cross-section of skin exposed (Fig. 2**d**). The slice was stained with a 1:100 dilution of Hoechst 33342 in *ex vivo* medium for 1 hour at 37 °C. After staining, the slice was placed on a poly-L-lysine (PLL)-coated dish with its cross-section side up (Fig. 2**f**), and the edges of the slice were fixed with agarose gel to maintain its position.

**Fig. 2.**
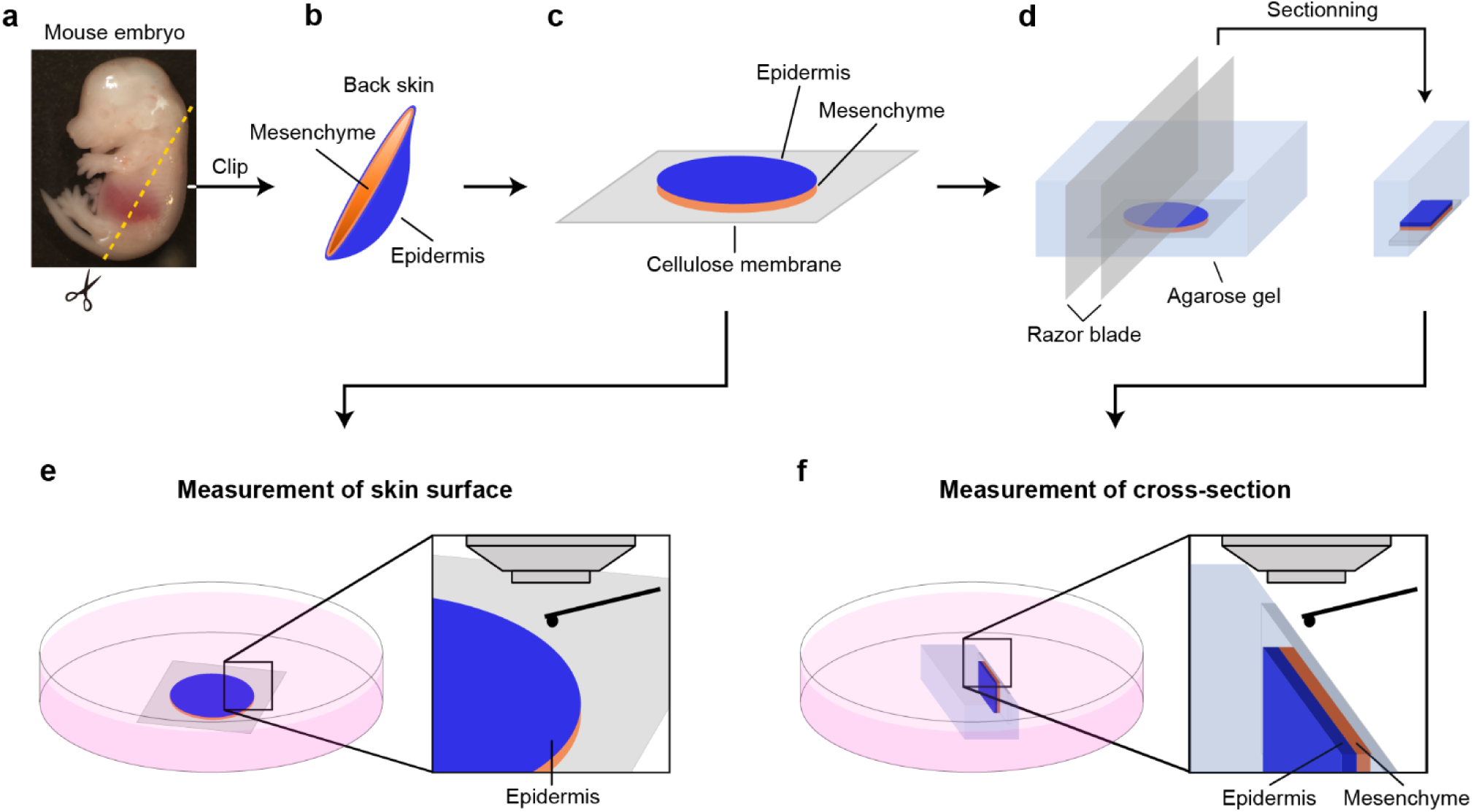
Preparation processes of the skin surface and slice samples. Surface sample preparation followed steps **a**–**c** and **e**; slice sample preparation followed steps **a**–**d** and **f**. **a**. Excision of skin from the back of a mouse embryo. **b**. Peeling of the skin tissue. **c**. Attachment of the tissue onto a cellulose membrane. **d**. Embedding of the tissue in agarose gel and stripping of the agar block. **e**. Surface measurement on the skin surface under *ex vivo* culture conditions. **f**. Cross-section measurement under *ex vivo* culture conditions.

### 2.4 Preparation of frozen-thawed slice samples

Cover glasses (MATSUNAMI, 24 × 24 mm, Thickness No. 1) were incubated in 0.01% PLL solution (Sigma, P4707) for 5 minutes and then air dried at room temperature. PLL-coated cover glasses (MATSUNAMI, C1210) were fixed to glass slides with mending tape. Dorsal tissues of E14.5 mice were excised, immediately embedded in optimal cutting temperature (O.C.T.) compound, and then frozen. The samples were then sealed and stored at −80 °C. Unfixed frozen samples were cut into thin slices (20 µm thick) and mounted on PLL-coated cover glasses. The slices were air dried at room temperature overnight. The edges of the cover glasses were surrounded with a Pap Pen (Asone, 1-5902-02) to create a repellent line. The tissue sections were washed three times with PBSto remove the O.C.T. compound. The slices were stained with a 1:100 dilution of Hoechst 33342 in the *ex vivo* culture medium for 1 hour at 4 °C. The cover glasses were removed from the glass slides and immersed in a 35 mm dish filled with the *ex vivo* culture medium.

### 2.5 Preparation of paraformaldehyde (PFA)-fixed slice samples

PFA (Wako, 163-20145) was dissolved in PBS (pH 7.4) at a final concentration of 4%. After preparing a slice sample using the described procedure (through Fig. 2**d** in Sec. 2.2), the slice was immediately fixed in 4% PFA fixative solution and immersed overnight at 4 °C. The following day, the slice was washed three times with PBS. Subsequently, the slice was stained with a 1:100 dilution of Hoechst 33342 in the *ex vivo* culture medium for 1 hour at 4 °C. After staining, the slice was placed on a PLL-coated dish with its cross-section side up (Fig. 2**f**), and the edges of the slice were fixed with agarose gel. The slice was stored at 4 °C until measurement.

### 2.6 Preparation of glyoxal-fixed slice samples

Glyoxal fixative solution was prepared according to a previous protocol [51]. Briefly, 28 ml of ddH_2_O, 7.89 ml of absolute ethanol, 3.13 ml of glyoxal (TCI, # G0152), and 0.3 ml of acetic acid (Wako) were vortexed thoroughly. The pH was adjusted to 4.0 with 1 N NaOH, and the solution volume was adjusted to 40 ml with ddH_2_O. The final concentration of glyoxal was 3%. After preparing (through Fig. 2**d** in Sec. 2.2) and immediately fixing a slice sample in 3% glyoxal fixative solution at pH 4.0, it was immersed overnight at 4 °C. The following day, the slice was washed three times with PBS. The slice was treated with NH_4_Cl (Wako, 013-02992) for 10 minutes to deactivate unreacted aldehydes, which might impede staining. Subsequently, the slice was stained with a 1:100 dilution of Hoechst 33342 in *ex vivo* medium for 1 hour at 4 °C. After staining, the slice was placed on a PLL-coated dish with its cross-section side up (Fig. 2**f**), and the edges of the slice were fixed with agarose gel. The slice was stored at 4 °C until measurement.

### 2.7 Enzyme treatment

Collagenase Type 4 (Worthington, LS004186), an enzyme that digests collagen, was added to slice samples at a concentration of 1 mg ml^-1^ for 1 hour under *ex vivo* culture conditions. AFM measurements and SIM observations were performed to compare the slices before and after treatment, and the region measured by AFM was traced using SIM images.

### 2.8 AFM measurement of tissue stiffness

Tipless silicon cantilevers (NANOSENSORS, TL-CONT) with borosilicate beads (Asone) were used for AFM measurments. The sizes of the beads were 24–26 µm in diameter for slice samples (Fig. 2**f**) and 81 µm in diameter for surface samples (Fig. 2**e**). The spring constants of the cantilevers were calibrated using a thermal noise method in water. Cantilevers with similar spring constants (0.2–0.4 N/m) were selected for the measurements. The applied force was 10–20 nN. The specific parameters for AFM measurement were a ramp size of 15 µm, a forward velocity of 50.0 µm/s, and a reverse velocity of 200.0 µm/s. Measurements were performed in *ex vivo* medium at 37 °C for living and frozen-thawed samples and in PBS at room temperature for PFA- and glyoxal-fixed samples. Force curves were obtained using contact mode force mapping. Fluorescence images were captured with a CMOS camera (Zeiss, Axiocam 712 mono) to determine the measured region. The obtained force curves were analyzed to calculate Young’s modulus by fitting the Hertzian model (spherical) using JPKSPM Data Processing software (Bruker).

### 2.9 Immunostaining and observation

Living slice and surface samples were immediately fixed in 4% PFA overnight at 4 °C. For immunohistochemistry, the samples were incubated in 0.5% Triton (MP Biomedicals)/PBS for 30 minutes for permeabilization and then incubated in CAS-Block™ Histochemical Reagent (Life Technologies, 008120) for 15 minutes for blocking, followed by washing with PBS. Subsequently, the samples were incubated with primary antibodies overnight at 20 °C and then incubated with secondary antibodies overnight at room temperature, followed by washing with PBS. The primary antibodies used in this study were anti-mouse Col I (1:200; Abcam, AB88147) and anti-rabbit ZO-1 (1:125; Invitrogen, 61-7300). The secondary antibodies were Goat Anti-Mouse IgG H&L Alexa Fluor 488 (1:200; Abcam, ab150113) and Goat Anti-Rabbit IgG H&L Alexa Fluor 488 (1:125; Abcam, ab150077). Hoechst 33342 was used to counter-stain nuclei, and Phalloidin-iFluor 555 (1:500; Abcam, ab176756) was used to stain F-actin. Stained samples were mounted with glycerol (Nacalai tesque) and then observed using confocal laser microscopy (Zeiss, LMS800).

### 2.10 Statistics and spatial profile analysis

To investigate the stiffness of the epidermis and mesenchyme on the skin cross-section, we manually extracted their regions from Hoechst-stained images. To assess the stiffness of the reticular and cell medial regions on the skin surface, we manually extracted their regions from Young’s modulus maps. During this process, we ignored the border regions between reticular or medial regions due to the large diameter of the beads attached to the cantilever. For statistical analysis, we calculated the average, standard deviation, and coefficient of variation to evaluate the stiffness values. In addition, to explore the spatial pattern of stiffness, we calculated the correlation coefficient of stiffness as a function of distance within each epidermis and mesenchyme. We determined the characteristic length of spatial correlation by fitting this coefficient function with an exponential function. The statistical values were derived from three or more samples.

ANOVA was conducted to determine whether a statistically significant trend occurred across the entire population. Tukey’s test was used to determine whether statistically significant differences existed between individual samples. All statistical analyses, including Tukey’s test and two-way ANOVA, were performed using EZR [52], which is a graphical user interface for R (The R Foundation for Statistical Computing, Austria). A confidence interval of 95% (corresponding to a P-value less than 0.05) was used to designate differences as statistically significant.

## 3. Results

### 3.1 USIM-AFM provides a clear paired fluorescence image and stiffness map of thick tissue slices, precisely coordinated at the subcellular scale

To investigate biological questions at the tissue scale, it is essential to distinguish the relevant cellular structures inside the target tissue during an observation. However, it becomes increasingly difficult to obtain a clear image using conventional inverted microscopes as the sample becomes thicker. To clearly observe the tissue surface and distinguish cellular structures, we combined USIM with AFM. We chose mouse embryonic skin, which consists of a layered epidermis and mesenchyme, as a representative sample with a spatially heterogeneous structure to validate the measurement principle.

To distinguish the heterogeneous structures of the layered epidermis and mesenchyme of mouse embryonic skin, we first tested the visibility of different illumination setups. The sample was sliced at 1–2 mm thickness and stained using a Hoechst dye. USIM clearly visualized the nuclear distribution in the top layer of the sliced mouse embryonic skin (Fig. 3**a**). We clearly distinguished the epidermis, with its densely packed, bright nuclei, from the mesenchyme, which had sparsely packed nuclei with darker signals (Fig. 3**a**). The magnified image even captured the shapes of individual nuclei densely distributed in the epidermis (Fig. 3**d**). The upright optical path, even without structured illumination, helped to distinguish the epidermis and the mesenchyme to a certain extent; these layers were visualized as regions with different brightnesses due to the differences in nuclear density inside (Fig. 3**b**). However, the shape of each nucleus was unclear in both the wide-field and magnified views (Fig. 3**e**). The conventional inverted fluorescence microscope was unable to detect either the individual nuclear shapes or the difference in nuclear density between the epidermis and the mesenchyme (Fig. 3**c, f**). These results demonstrate a real advantage of adopting USIM to recognize the structure in the top layer of a thick tissue slice.

**Fig. 3.**
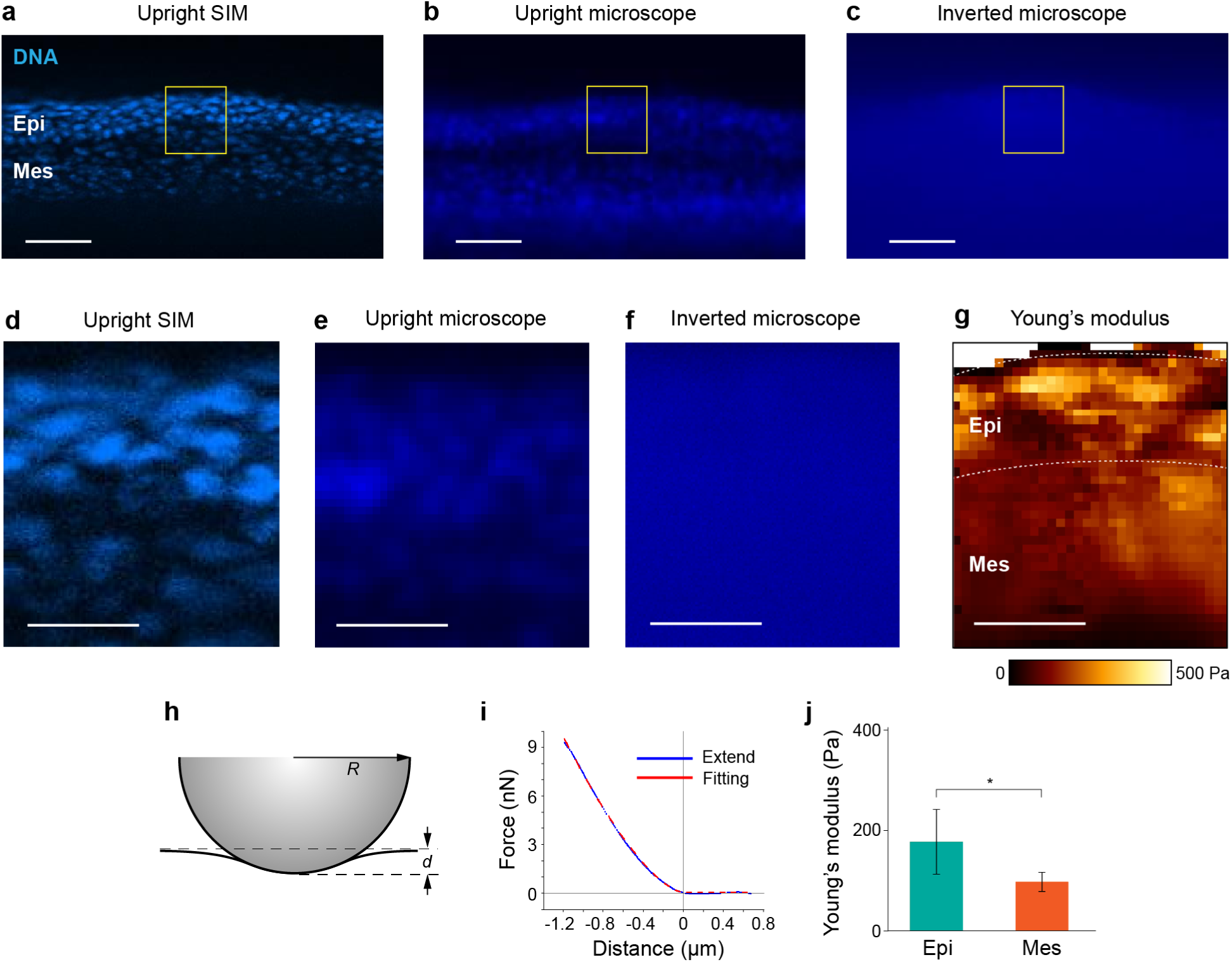
Stiffness and fluorescence maps on the same focal plane of a living skin slice. **a**–**c** Fluorescence images on the top side of a living skin slice acquired by upright structured illumination microscopy (**a**), conventional upright microscopy (**b**), and inverted fluorescence microscopy (**c**). **d**–**f**. Enlarged images of macroscopic views (**a**–**c**). **g**. Stiffness map of a living skin slice on the same focal plane with the fluorescence images (**d**–**f**). White dotted lines indicate the boundaries of the epidermis and mesenchyme. **h**. Schematic of the contact point between an atomic force microscopy (AFM) probe and the sample surface. The probe has a spherical tip of radius *R*. The amount of indentation is *d*. **i**. Typical force curve obtained by the measurement. The blue curve indicates a force curve when the AFM probe is pushed into a sample. The red dotted curve indicates a force curve fitted by the Hertzian model. **j**. Elastic modulus value of the epidermis and mesenchyme. Data represent the mean ± s.d. (N = 4). *P < 0.05 (Welch’s t-test). Scale bars, 50 μm (**a**–**c**); 20 μm (**d**–**g**).

AFM measurement was then carried out to obtain the stiffness map of the top layer of the tissue slice at the corresponding site. Based on the live observation using USIM, we selected a region that contained the boundary between the epidermis and the mesenchyme, precisely setting the border at the middle position of the measurement field of the atomic force microscope. The stiffness in this region was then mapped using an atomic force microscope equipped with a tipless cantilever with a bead (Fig. 3**g**). The indentation, represented by *d*, of approximately 3–4 µm, and the bead radius, represented by *R*, of approximately 24–26 µm, characterize the force-distance measurement (Fig. 3**h**). The obtained force curves perfectly fit to the Hertzian model in the range of *d*/*R* < 20% (Fig. 3**i**); this agreement ensured that the sliced tissue was sufficiently thick to exhibit linear elasticity in this range, *i.e.*, the mechanical characteristics remain unaffected by environmental factors such as the stiffness of the agar and the supporting membrane [1]. Young’s modulus was then calculated based on this force-distance relationship, setting Poisson’s ratio to 0.5, as commonly used in soft tissue mechanics [53]. Sequential AFM measurements in the field of view provided a stiffness map around the boundary, clearly visualizing a difference in stiffness distribution between the epidermis and mesenchyme (Fig. 3**g**). The epidermis showed a highly heterogeneous stiffness distribution containing stiffer islands at approximately 10 µm. In contrast, the mesenchyme showed a relatively homogeneous stiffness distribution at a considerably lower value. Statistical comparison also confirmed a significant difference in the mean Young’s modulus between the epidermis (177 ± 64 Pa) and the mesenchyme (98 ± 19 Pa) (Fig. 3**j**). Notably, the difference in Young’s modulus corresponded to that in the nuclear density as well as the signal intensity (Fig. 3**d**).

The stiffness of live embryonic skin has not been investigated despite its importance in tissue morphogenesis during development. The absolute stiffness value of mouse embryonic skin measured using our USIM-AFM method was 100–200 Pa, significantly softer than human skin tissues, which have stiffness values of approximately 70–160 MPa measured by uniaxial tensile tests [54]. Conversely, these measured stiffness values are of the same order as those measured for mouse embryonic stem cell-derived retinal organoids at approximately day 9, corresponding to E11.5, *e.g.*, neuroepithelium (321 ± 89 Pa), neural retina (146 ± 44 Pa), and retinal pigment epithelium (391 ± 124 Pa) [39], supporting the validity of our measurement. This finding also highlights a stiffness difference between the epidermis and mesenchyme at the microscopic scale, correlating with the cell density inside. This observation is consistent with conventional studies reporting that the tissue stiffness increases together with the cell density during mesodermal convergent extension in gastrulation [55], enhanced neurogenesis in adult mice [56], and the later stages of brain development in Xenopus embryos [40]. Our new measurement system enabled the successful characterization of mechanical properties in soft skin tissue during development with microscopic precision. The integration of USIM with AFM was particularly effective in precisely identifying tissue structures and in acquiring spatially coordinated stiffness maps and fluorescence images of thick live tissues. We thus utilized this USIM-AFM method to explore the tissue-scale mechanics of mouse embryonic skin in the following sections.

### 3.2 Stiffness distribution detected on the intact embryonic skin surface reveals heterogeneous properties characterizing epithelial junctions and epidermal cell bodies

The USIM-AFM method enabled us to measure the absolute stiffness value and its spatial distribution on the top layer of a live tissue section. Focusing on tissue-scale functionality and integrity, the technique remains invasive, as it requires slicing a sample from an embryonic body. The necessity of such an invasive process to obtain accurate *in vivo* stiffness values remains an open question. Next, to clarify the importance of tissue sectioning, we measured the stiffness of an intact epidermis noninvasively without slicing and compared these measurements to those from sliced tissue sections.

The Young’s modulus value of the entire intact skin surface was approximately 217 ± 21 Pa, a value that was unexpectedly higher than that measured for the epidermis of the sliced cross-section, which was approximately 177 ± 64 Pa (Fig. 3**j**, 4**b**). Furthermore, the stiffness map of the intact epidermis revealed a distinct reticular pattern (Fig. 4**a**). By isolating the reticular and medial regions, we were able to calculate the stiffness in each of these regions (Fig. 4**c**). The value of Young’s modulus in the medial region was 183 ± 14 Pa, which was significantly lower than the value of 243 ± 25 Pa found along the reticular pattern (Fig. 4**b**). This value was comparable to that measured on the epidermis of the sliced cross-section. We hypothesized that this unique pattern of stiffness distribution on the intact skin arose from the tissue structure of the epithelium. Thus, we stained ZO-1, a scaffold protein recruited in mature tight junctions, using a fixed sample. Fluorescence observation showed ZO-1 distributed in a reticular pattern (Fig. 4**d**–**f**), mirroring the reticular geometry and its size in the stiffness map (Fig. 4**a**). These results show that the reticular region, where higher stiffness was detected, corresponds to epithelial junctions, indicating that the stiffness of the medial region reflects that of epidermal cell bodies.

**Fig. 4.**
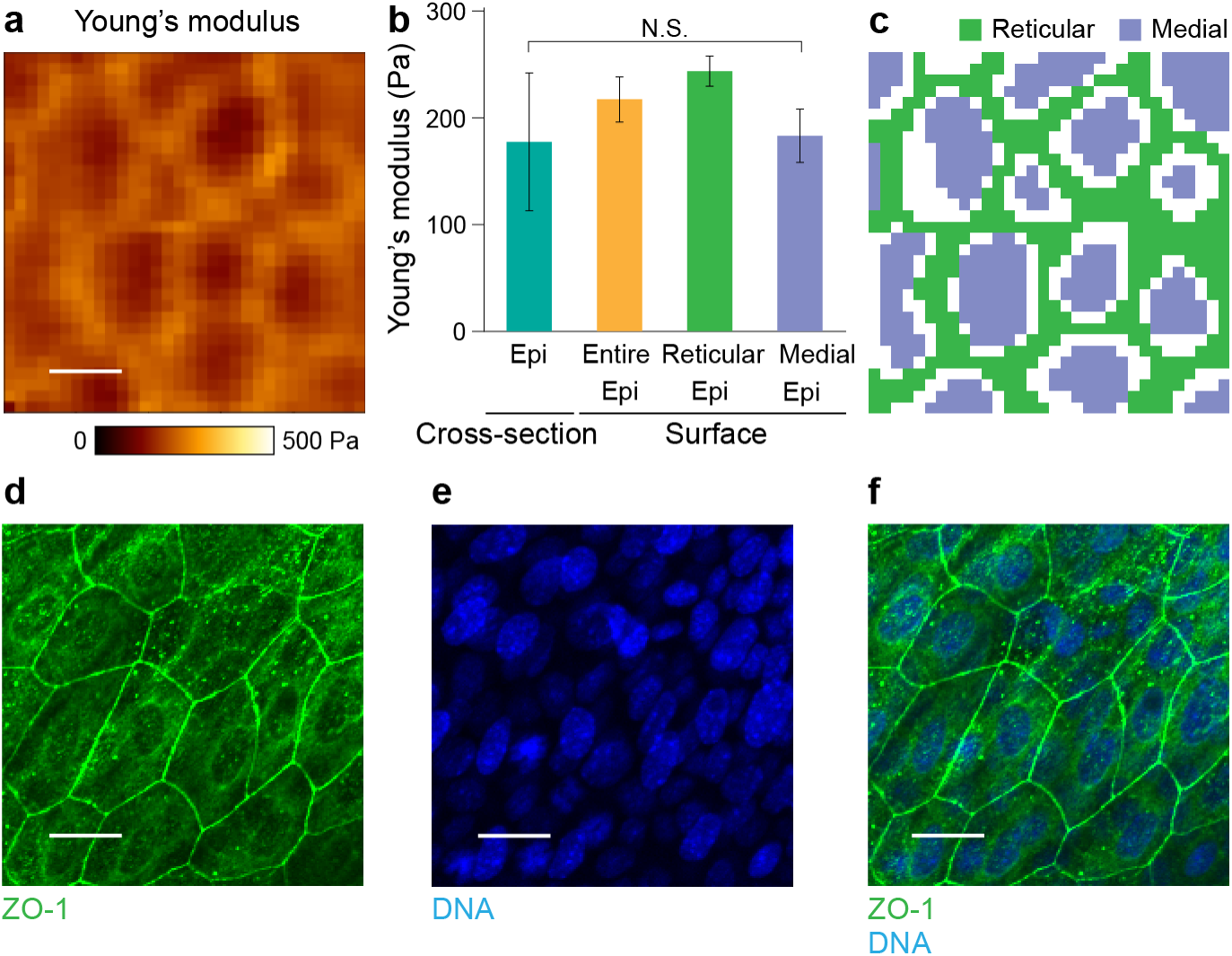
Validation of the living tissue slice measurements by comparison with surface measurements. **a**. Stiffness map of the living skin surface. **b**. Comparison of elastic modulus values from the epidermis region of the cross-section, the entire surface region, and the reticular and medial regions on the surface. Data represent the mean ± s.d. (N = 4). *P < 0.05, as determined by Tukey’s test. **c**. Masked image highlighting the reticular pattern. In the masked image, the purple region indicates the cell medial region, and the green region denotes the reticular region **d**–**f**. Immunostained images of the skin surface sample showing ZO-1 (**d**), DNA stained with Hoechst (**e**), and their superimposition (**f**). Scale bars, 20 μm (**a**, **d**– **f**).

Mature epithelial junctions form only at the embryonic skin surface, not within the epidermis [57]. Our comparison revealed that the average stiffness measured on the intact skin surface was highly influenced by these junctions, particularly the stiffness detected in the reticular patterns. Conversely, the stiffness of the medial region, despite being measured at the skin surface, corresponded to that of the underlying epidermis. This quantitative comparison demonstrates that the stiffness distribution of the intact skin surface reflects diverse information from cellular and tissue structures, highlighting the importance of subcellular resolution in measurements. Simultaneously, this result emphasizes the need for caution when estimating tissue mechanical properties from limited surface information. This observation partially justifies the need for tissue sectioning to accurately measure the stiffness of 3D tissues deep within the body.

### 3.3 Spatiotemporal observation of collagenase-treated tissue slices reveals the role of collagen as a principal mechanical component of embryonic skin in early development

AFM has limited flexibility in the optical observation of the sample surface during measurement due to the contact-based principle. This limitation is particularly manifested when handling a thick, optically opaque sample; common inverted microscopes are not applicable for observing the top surface of such thick samples. Thus, AFM measurement and fluorescence observations need to be separately carried out. However, the process of transferring live, fragile tissue across microscopes can damage the sample. Additionally, such iterative movements of the sample are not suitable for time-lapse observation, which is one of the core interests of live measurement. Our USIM-AFM method, which enables simultaneous fluorescence observation and mechanical characterization of a thick tissue sample, solves this problem and perfectly fits time-lapse observation. Leveraging this advantage, we demonstrate the time-course measurement of live tissue slice stiffness.

Collagen, a major mechanical component sustaining the tissue structure, has been recently shown to dynamically remodel during development [58], thus playing a significant role in tissue morphogenesis. However, its quantitative contribution to the stiffness of developing tissues remains unclear. Utilizing the USIM-AFM method, we conducted a time-course measurement of live tissue slice stiffness in collagenase-treated tissues to reveal the contribution of collagen to the stiffness of developing embryonic skin. Prior to collagenase treatment, we first observed immunostaining for collagen I, the actin cytoskeleton, and nuclei in a fixed slice sample (Fig. 5**a–d**). Collagen I and the actin cytoskeleton were strongly localized at the junctional parts in the epidermis, while these molecules were distributed more homogeneously in the mesenchyme (Fig. 5**a**). The fluorescence intensities of collagen I and nuclei were higher in the epidermis, confirming a clear difference in subcellular structures between the epidermis and mesenchyme. This observation of collagen I is consistent with those in a previous study, although the results are not explicitly mentioned [59].

**Fig. 5:**
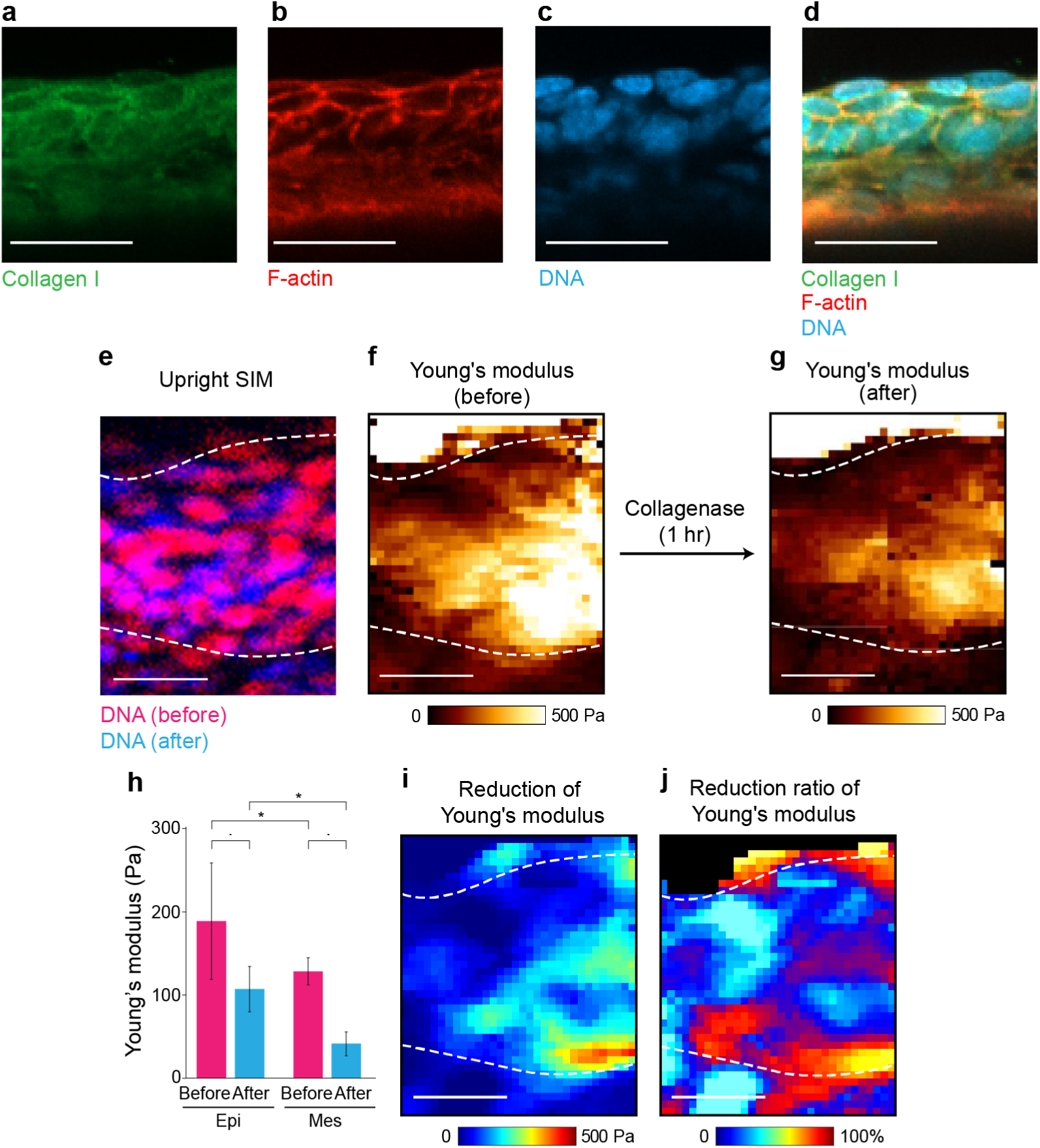
Quantification of temporal changes in stiffness of living tissue slices. **a**–**d.** Immunostaining of a slice sample. Collagen I (**a**); F-actin stained with phalloidin (**b**); DNA stained with Hoechst (**c**); superimposition of **a**–**c** (**d**). **e.** DNA images of a slice sample before and after the addition of Collagenase Type 4. **f**, **g**. Stiffness maps before and 1 hour after the addition of Collagenase Type 4, respectively. **h, j.** Difference and quotient of the stiffness maps in **f** and **g**. **j**. Average stiffness of the epidermis and mesenchyme before and 1 hour after the addition of Collagenase Type 4. Data represent the mean ± s.d. (N = 3). *P < 0.05, as determined by two-way ANOVA followed by Tukey’s test. Scale bars, 50 μm (**a**–**d**); 20 μm (**e**-**i**).

We conducted simultaneous live measurements of sliced tissue using USIM-AFM. We first set the field of view containing both the epidermis and the mesenchyme based on the USIM observation of nuclei vividly stained with Hoechst dye. To quantitatively analyze the extent to which collagen contributes to the mechanical strength of embryonic skin, we added Collagenase Type 4, an enzyme that digests a wide range of collagens, to a live tissue slice and tracked the stiffness distribution using AFM. Nuclei were also observed using USIM both before and after the enzymatic treatment (Fig. 5**e**). The tissue inevitably deforms during collagenase treatment as the digestion of collagens releases the residual stress retained within the extracellular matrix. We first compared the nuclear positions before and after the enzymatic treatment to assess the displacement field within the tissue (Fig. 5**e**). Using the nuclear positions as references, we then obtained corrected stiffness maps before and after the treatment, aligning them with each other (Fig. 5**f, g**). The enzymatic treatment considerably reduced the stiffness of both the epidermis and mesenchyme (Fig. 5**h**). Quantitatively, Young’s modulus of the epidermis decreased from 189 ± 70 Pa to 128 ± 27 Pa and the mesenchyme decreased from 107 ± 16 Pa to 41 ± 14 Pa. The stiffness reduction rate was much greater in the mesenchyme (61%) than in the epidermis (32%). We then focused on the spatial variance in the stiffness change (Fig. 5**i**, **j**). The decrease in the absolute value of Young’s modulus spatially correlated well with the initial stiffness (Fig. 5**f, i**). In contrast, the relative reduction, *i.e.*, the stiffness change normalized by the initial stiffness value, was observed to be more homogenous across the entire tissue (Fig. 5**j**). These observations indicated that the collagenase uniformly digested the collagens across the epidermis and the mesenchyme (Fig. 5**a**, **d**). Furthermore, the fact that the collagenase treatment nearly halved the absolute stiffness value of both tissues suggests that collagen is the primary mechanical contributor to skin stiffness, even during development. These results demonstrate that the simultaneous acquisition of spatially coordinated fluorescence images and stiffness maps using USIM-AFM offers significant advantages in the live measurement of thick tissue, a capability that is fundamentally challenging for conventional AFM to provide.

### 3.4 Live measurement without preservation treatment is crucial to understand tissue mechanics at a subcellular resolution

Tissue-scale mechanical testing is often carried out using samples preserved in various ways [41,43,46,47,57,60]. This practice arises due to practical limitations in sample storage and transport. However, recent studies have revealed that these preservation processes critically influence the stiffness values measured by AFM [41]. Therefore, we utilized our USIM-AFM method to gain a deeper understanding of how preservation processes alter the mechanical properties of tissue samples at the microscopic level and, in turn, to clarify the extent to which live measurement is important in tissue-scale mechanical characterization. We prepared three types of treated samples, frozen-thawed samples without chemical treatment, samples fixed with 4% PFA, and samples fixed with 3% glyoxal, as used in a previous study [41]. We quantitatively compared the stiffness maps of these treated samples with those from living samples.

The samples were first stained with Hoechst dye, as used for the living samples described previously (Sec. 3.1), to visualize the nuclear distribution inside. Based on the Hoechst fluorescence observed with USIM, we distinguished the border between the epidermis and the mesenchyme and set the field of view containing the layered tissue structure (Fig. 6**a–d**). The stiffness map at the center of the USIM image was obtained using AFM (Fig. 6**e–h**). The range of absolute stiffness values detected in the treated samples (Fig. 6**f–h**) was generally approximately an order of magnitude higher than that detected in the living sample (Fig. 6**e**). Note that Fig. 6**e–h** are colored based on scales of different ranges. However, it should be noted that the relative difference in stiffness between the epidermis and the mesenchyme was still clearly exhibited in all treated samples. Beyond the absolute stiffness values, the texture of the obtained stiffness maps also appeared different. Frozen-thawed samples showed spatially smoother stiffness distributions (Fig. 6**f**) than those of living samples (Fig. 6**e**). In contrast, PFA- and glyoxal-treated samples showed rougher stiffness distributions (Fig. 6**g, h**). We further calculated spatial correlation functions for each stiffness map to quantitatively assess the impact of the fixation processes on the stiffness pattern (Fig. 6**i–l**). The correlation distance, being the decay constant of the curve, characterizes the representative scale of the texture pattern in the image.

**Fig. 6:**
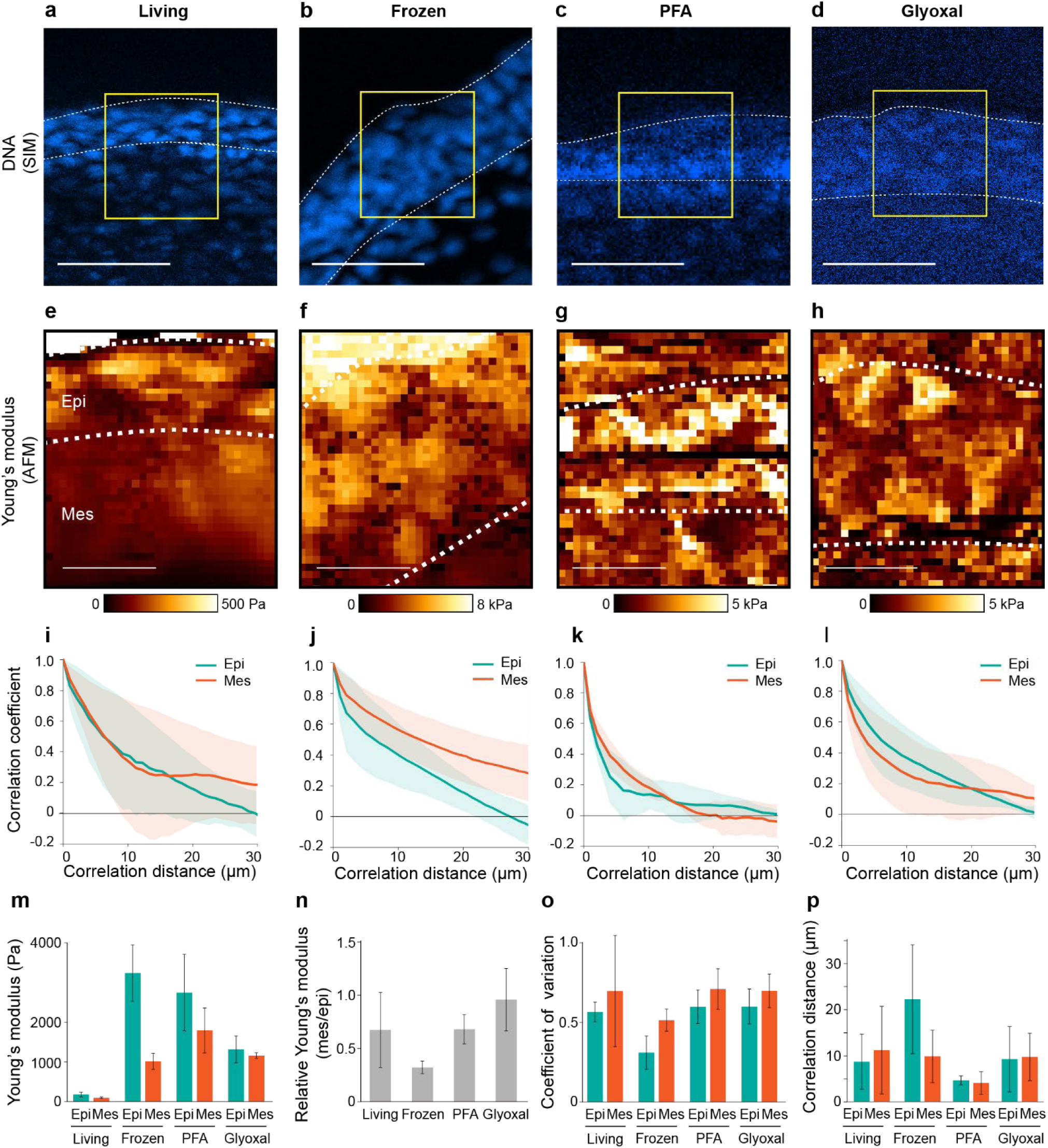
Comparison of stiffness across living, frozen-thawed, paraformaldehyde (PFA)-fixed, and glyoxal-fixed samples. **a**–**d,** DNA stained with Hoechst. White dotted lines indicate the boundaries of the epidermis. Yellow squares indicate the regions measured by atomic force microscopy. **e**–**h,** Stiffness maps of each selected region in **a**–**d**. **i**–**l**, Spatial correlation coefficient of stiffness as a function of distance. Blue and orange lines indicate the epidermis and mesenchyme. **m.** Average stiffness in the epidermis and mesenchyme regions. **n.** Relative stiffness of the epidermis and mesenchyme regions. **o**. Variation coefficient of stiffness in the epidermis and mesenchyme regions. **p**. Correlation distance of stiffness in the epidermis and mesenchyme regions. Data represent the mean ± s.d. (n = 4 for living, n = 6 for frozen-thawed, n = 3 for PFA-fixed, and n = 3 for glyoxal-fixed samples). Scale bars: 20 μm (**a**–**h**).

Quantitative comparison of the mean stiffness values of the epidermis and mesenchyme showed that all fixation processes increased the absolute stiffness values 10 to 20 times compared with those of living tissue (Fig. 6**m**). Focusing on the relative mesenchyme to epidermis stiffness ratio, which is approximately 0.7 in living tissue, it is evident that these fixation processes had different impacts on the epidermis and mesenchyme (Fig. 6**n**). While the freeze-thaw process expanded the relative difference, glyoxal fixation tended to reduce this difference. Notably, PFA fixation maintained the mesenchyme/epidermis stiffness ratio at approximately 0.7. However, the relative mesenchyme/epidermis stiffness ratio generally remained at similar values below 1, indicating the magnitude relationship of stiffness between the epidermis and mesenchyme is maintained throughout the preservation processes to a certain extent. This finding is supported by the fact that the coefficient of variation, an indicator of stiffness deviation normalized across the entire image, for the four AFM images remained at similar values near 0.5, except for the frozen-thaw epidermis (Fig. 6**o**). These indicators suggest that the magnitude difference in tissue stiffness remains at regional scales, such as the epidermis, mesenchyme, or the entire field of view, throughout the preservation process. Nevertheless, the stiffness maps of these samples suggested different textures (Fig. 6**e–h**). In fact, the correlation distance was larger in the frozen-thaw epidermis, while it was generally smaller both in the epidermis and the mesenchyme of the PFA-fixed samples (Fig. 6**p**). These differences in the correlation length clearly indicate that microscopic features of the stiffness pattern, which likely derive from subcellular structures, are impaired by these preservation processes. However, there was no considerable difference in the correlation distance of the stiffness distribution between the live and glyoxal-fixed samples, implying that glyoxal fixation exerts a modest influence on the microscopic features of the stiffness distribution.

These quantitative comparisons revealed the different impacts of preservation processes on tissue samples, as summarized in Table 1. Freeze-thaw, PFA fixation, and glyoxal fixation all preserved relative differences in stiffness between the epithelium and mesenchyme; however, none of these techniques maintained the original stiffness values measured in live tissue. The freeze-thaw process tended to increase the relative stiffness difference between tissues. It also flattened the stiffness distribution. The PFA fixation process exerted the opposite influence, making the stiffness distribution rougher, yet it effectively maintained the mesenchyme to epidermis stiffness ratio. Glyoxal fixation tended to decrease the relative tissue difference but had the mildest impact on the microscopic features of the stiffness distribution among these fixation processes. These results may indicate that treated samples are still useful for evaluating the relative difference across various tissues, *i.e.*, epidermis and mesenchyme. However, the microscopic features of the stiffness distribution at the subcellular scale are generally deteriorated by preservation processes. Although the live stiffness measurement of tissue-scale samples involves technical complications, there is currently no alternative method to measure the absolute stiffness value or to capture the spatial pattern at the subcellular scale, which is comparable to microscopic images commonly used in molecular biology.

**Table 1:** Applicability of sample preparation to stiffness measurement.

|  | Living | Freeze-thaw | PFA* | Glyoxal |
| --- | --- | --- | --- | --- |
| Relative average difference between various types of tissues | ✓ | ✓ | ✓ |  |
| Statistical variation in each type of tissue | ✓ |  | ✓ | ✓ |
| Spatial correlation in each type of tissue | ✓ |  |  | ✓ |

\*PFA, paraformaldehyde

## 4. Discussion

In this study, we proposed a new technique that integrates AFM, USIM, and *ex vivo* slice culture to evaluate stiffness in spatial coordination with molecular distributions in 3D tissue (Fig. 1). A thick slice of living mouse embryonic skin, a model tissue consisting of mechanically heterogeneous layers, was chosen to demonstrate the measurement principle (Fig. 2). Because of the visibility provided by USIM, we successfully distinguished the layered epidermis and mesenchyme of sliced living skin and obtained a spatially coordinated stiffness map and fluorescence image (Fig. 3). This quantitative and precise approach enabled us to observe the highly heterogeneous stiffness distribution of the intact embryonic skin surface (Fig. 4), revealing that mature epithelial junctions, found only on the top skin surface, critically increase the tissue stiffness. This observation, in turn, underlines the importance of *ex vivo* slice culture as a realistic compromise to determine the mechanical properties of living tissue located deep inside a body. The USIM-AFM method also allowed us to track the time course of stiffness changes in embryonic skin treated with collagenase (Fig. 5). The precisely aligned stiffness maps and nuclear images of collagenase-treated tissue slices at different time points quantitatively demonstrated that collagens are the major mechanical components endowing stiffness to the embryonic skin. Furthermore, USIM-AFM images revealed that preservation processes, often involved in mechanical characterization of tissue, alter both the stiffness value and its distribution at microscopic resolution (Fig. 6). To the best of our knowledge, this study represents the first attempt to thoroughly explore live spatiotemporal mechanics maps of thick tissue slices in combination with microscopic observation.

The series of USIM-AFM observations of living embryonic skin slices revealed that the stiffness distribution within the skin is highly heterogeneous in three dimensions. The stiffness detected on the intact skin surface was locally enhanced by mature epithelial junctions, resulting in a reticular pattern with an increased Young’s modulus (Fig. 4). In the deeper part of the slice, the epidermis exhibited generally greater stiffness than did the mesenchyme, and there were distinct patterns of stiffer and softer regions within each tissue type (Fig. 3). However, the causes of such spatial heterogeneity in stiffness within each tissue type remain unclear. This finding may be attributed to the internal tissue structures, such as hair placodes and vessels [61]. Additionally, intracellular structures, including condensed chromatin in the nucleus, are likely to contribute to such spatial heterogeneity, considering the correlation of the tissue stiffness and Hoechst signal intensity in these tissues (Fig. 3). Utilization of GFP knock-in mice for live observation of labeled candidate molecules, together with AFM measurements, would further clarify the role of individual molecules in the formation of such a mechanically heterogeneous environment inside living tissue. When combined with genetic manipulation tools, USIM-AFM could become a powerful approach to quantitatively understand the causal relationship between the expression levels of specific target molecules and tissue mechanical heterogeneity with subcellular resolution.

Frozen-thawed or chemically fixed samples are commonly used in AFM measurements to estimate tissue mechanical properties, as referenced in several studies [41,43,46,47,60]. However, the extent to which stiffness values measured in treated tissues reflect the genuine stiffness of living tissue has not been adequately verified or discussed in many studies, except for a few pioneering works reporting a hardening effect of glyoxal fixation [41]. Our results distinctly demonstrate, for the first time, that frozen-thawed, PFA-fixed, and glyoxal-fixed samples exhibit a significantly altered tissue stiffness pattern compared with that of *ex vivo* maintained living samples (Fig. 6). Specifically, these treatments increased the average Young’s modulus of the tissue by approximately 10-fold (Fig. 6**m**). Additionally, these treatments influenced the stiffness distribution differently, either flattening or roughening it at the subcellular scale (Fig. 6**p**). Such a microscopic difference may indicate that the stiffness changes caused by these treatments are attributed to distinct reasons. For example, repeated dehydration induced by freeze-thawing denatures and condenses the biomolecules contained in the sample, resulting in significant changes in tissue stiffness. Conversely, PFA, a common fixative, crosslinks biomolecules almost non-selectively, resulting in the formation of large insoluble biomolecule condensates [48]. Glyoxal, similar to PFA, is an aldehyde-based chemical cross-linker known to alter collagen stiffness [62]. Chemical crosslinks formed by these treatments are likely to increase tissue stiffness by limiting the deformability of the biomolecules. Because stiffness is a simple mechanical parameter that can reflect all properties of the molecules contained inside, it is difficult to clarify the mechanism of how the tissue stiffness is altered by each treatment. It is, however, notable that the magnitude relationship of the stiffness between the epidermis and mesenchyme tends to be maintained at the regional scale, regardless of the treatment type (Fig. 6**n**, **o**). This observation is consistent with a previous study reporting that glyoxal fixation increased the absolute stiffness of brain tissue by approximately 3-fold, while the magnitude relationships between the subventricular zone, ventricular zone, intermediate zone, and cortical plate were maintained even after fixation [41]. We thus conclude that preserved samples remain useful for assessing the relative differences between various tissues. However, the effects of freezing or fixation with PFA and glyoxal on tissue stiffness should not be overlooked if we aim to find meaning in the absolute stiffness values as well as their distribution with subcellular resolution. Our observations suggest that the capability of live tissue stiffness measurement provided by our *ex vivo* USIM-AFM system offers significant insights beyond the complexity of the instrumentation involved.

Our USIM-AFM observations revealed that the tissue stiffness close to the body surface is highly heterogeneous in three dimensions (Fig. 4). This finding highlights the importance of comparing high-resolution stiffness maps from different directions, such as the body surface and tissue cross-sections, to fully understand the mechanical properties of 3D living tissue. Despite the significant potential of USIM-AFM, there is room for improvement in increasing the 3D resolution of stiffness distribution, particularly the notable gap in *z*-resolution between stiffness maps and fluorescence observations. Specifically, the z-resolution of stiffness mapping, determined by the indentation depth, is approximately a few microns in macroscopic measurements. Conversely, the current finest *z*-resolution of USIM, determined by the numerical aperture of the objective lens and the wavelength of light, is approximately 5 µm. The inadequate *z*-resolution of optical imaging, compared to the mechanical characterization, makes it difficult for the current setup to precisely determine whether structures seen in fluorescence images are closer to the top or bottom of the plane where AFM detects stiffness. This issue could be addressed by integrating upright confocal microscopy with AFM using custom configurations. Furthermore, application of nano-endoscopy AFM, which measures intracellular structures using penetrating needle probes [63], could significantly enhance the resolution of 3D mechanical characterization of tissue. By combining improvements in optics and mechanical probes, USIM-AFM could provide a novel methodology to comprehensively characterize the 3D mechanical properties and molecular distributions at micron resolution within living tissue specimens.

Currently, there exists a trade-off between reliability and non-invasiveness in the mechanical characterization of tissue-scale samples. In this regard, AFM-based techniques and optical elastography complement each other. In principle, AFM operates as a contact-based scanning probe microscopy technique. Therefore, the sample must be sectioned to expose the deep-seated region of interest to measure local stiffness (Fig. 4**b**). Accordingly, while our USIM-AFM method maintains the living tissue sample through *ex vivo* slice culture, it is inevitably invasive. Despite its invasiveness, USIM-AFM offers highly reliable absolute stiffness values of tissue samples, grounded in a precise force-distance relationship (Fig. 3**i**). This reliable characteristic of AFM, not limited to targets being linearly elastic, ensures the broad applicability of USIM-AFM across various biological samples. For example, mechanical testing even under large deformation can also be potentially conducted with a living tissue slice if an appropriate model is applied. Conversely, Brillouin microscopy, an advanced form of optical elastography, enables 3D mechanical property measurement without the need for sectioning. This is a profound merit that AFM-based techniques have a fundamental difficulty providing. However, the Brillouin signal, which is a frequency shift caused by the mechanical property of the target, is also sensitive to the state of the matter, including amorphousness and crystallinity. Consequently, correlating the Brillouin signal with the actual mechanical properties of the target remains challenging, necessitating careful calibration, especially with biologically heterogeneous samples. Our USIM-AFM technique could offer a robust and precise reference for absolute stiffness to the evolving field of optical elastography. Thus, USIM-AFM and optical elastography may emerge as complementary tools, mutually enhancing the exploration of soft tissue mechanobiology.

Since the discovery of stiffness-dependent differentiation of mesenchymal stem cells [3], the Young’s modulus value of the extracellular environment and its impact on cell fate have been core interests in the field of mechanobiology. Environmental stiffness is closely linked to the composition of the extracellular matrix (ECM) and is dynamically altered by various cellular activities [64]. Nevertheless, its mechanobiological role in living tissues has not been extensively explored in the long term. Our USIM-AFM method will be a prominent tool to reveal the causal relationship between tissue stiffness and the functionality of biological molecules, especially if combined with genetic tools that vividly label the target molecule. This approach can also be applied to assess the *in vivo* therapeutic effect of recently developed mechanotherapies [65] by quantitatively comparing the mechanical effects and the biological functionality of the target tissue. Because the impact of such mechanical cues from the extracellular environment is known to be memorized by cells [66], mechanical influences have also been actively incorporated into the design of biomaterials that aim to guide cellular fates, such as differentiation [67]. Mechanical properties such as stiffness and viscosity are, undoubtedly, critical parameters in designing cutting-edge biomaterials. As mechanobiological questions become more closely aligned with *in vivo* events, *in vitro* tissue culture platforms have increased in complexity to capture cell-cell and cell-ECM interactions in three dimensions. The USIM-AFM method could also contribute to evaluating the interaction between mechanically designed biomaterials and cellular behaviors in engineered platforms. Our USIM-AFM method could serve as a technical basis to unveil the mechanobiological principles underlying various cellular behaviors inside 3D tissues by providing a solid methodology to explore the link between cellular mechanics and biomolecular distributions in tissue-scale vastness with micron-scale precision.

## 5. Conclusions

Combining AFM, SIM, and *ex vivo* slice culture, we have proposed a new technique, USIM-AFM, to obtain spatially coordinated stiffness maps and fluorescence images of living tissue slices. This technique offers a significant advantage in the spatiotemporal assessment of living tissue stiffness. A series of experiments using USIM-AFM demonstrated the importance of measuring the absolute stiffness value of live tissue sections to accurately understand the actual mechanical properties of tissues *in vivo*. Notably, quantitative comparison of stiffness distributions unveiled the distinct impacts of conventional tissue preservation techniques on tissue stiffness as well as its texture. Our USIM-AFM technique provides a solid methodology to obtain high-resolution stiffness maps that are comparable to biomolecular distributions, thus this technique could be a powerful tool to pursue mechanobiological questions in living 3D tissues.

## Acknowledgments

We would like to thank Dr. Kazuko Okamoto at Hiroshima University for discussions. This work was supported by the WISE program for Nano-Precision Medicine, Science, and Technology of Kanazawa University by the Ministry of Education, Culture, Sports, Science and Technology (MEXT); the Japan Science and Technology Agency (JST), CREST [Grant No. JPMJCR1921, JPMJCR1926]; the Japan Agency for Medical Research and Development (AMED) [Grant No. 21bm0704065h0003]; the Japan Society for the Promotion of Science (JSPS), KAKENHI [Grant Nos. 21H01209, 21KK0134, 22K18749, and 22H05170]; and the World Premier International Research Center Initiative, MEXT, Japan.

## Author Contribution Statement

IS prepared the samples, performed measurements, analyzed data, and wrote the first draft of the manuscript; RM prepared frozen-thawed slice samples; SO conceived of the project; RM, DW, and HF provided crucial ideas; IS and SO developed the microscope setup; IS, RY, TY, and SO wrote the manuscript. All authors contributed to the final manuscript.

## Data Availability Statement

The datasets analyzed during the current study are partially available from the corresponding author upon reasonable request.

## Notes

### Competing Interest Statement

The authors have declared no competing interest.

